# Analysis of community connectivity in spatial transcriptomics data

**DOI:** 10.1101/2022.04.11.487907

**Authors:** Carter Allen, Kyeong Joo Jung, Yuzhou Chang, Qin Ma, Dongjun Chung

**Author notes:** These authors contributed equally to this work (Carter Allen, Kyeong Joo Jung).

## Abstract

The advent of high throughput spatial transcriptomics (HST) has allowed for unprecedented characterization of spatially distinct cell communities within a tissue sample. While a wide range of computational tools exist for detecting cell communities in HST data, none allow for characterization of community connectivity, i.e., the relative similarity of cells within and between found communities – an analysis task that can elucidate cellular dynamics in important settings such as the tumor microenvironment. To address this gap, we introduce the concept of analysis of community connectivity (ACC), which entails not only labeling distinct cell communities within a tissue sample, but understanding the relative similarity of cells within and between communities. We develop a Bayesian multi-layer network model called BANYAN for integration of spatial and gene expression information to achieve ACC. We use BANYAN to implement ACC in invasive ductal carcinoma, and uncover distinct community structure relevant to the interaction of cell types within the tumor microenvironment. Next, we show how ACC can help clarify ambiguous annotations in a human white adipose tissue sample. Finally, we demonstrate BANYAN’s ability to recover community connectivity structure via a simulation study based on real sagittal mouse brain HST data.

**Availability:** An `R` package `banyan` is available at https://github.com/carter-allen/banyan.

**Supplementary information:** Supplementary data are available online.

**Author Summary:** The proliferation of spatial transcriptomics technologies have prompted the development of numerous statistical models for characterizing the makeup of a tissue sample in terms of distinct cell sub-populations. However, existing methods regard inferred sub-populations as static entities and do not offer any ability to discover the relative similarity of cells within and between communities, thereby obfuscating the true interactive nature of cells in a tissue sample. We develop BANYAN: a statistical model for implementing analysis of community connectivity (ACC), i.e., the process of inferring the similarity of cells within and between cell sub-populations. We demonstrate the utility of ACC through the analysis of a publicly available breast cancer data set, which revealed distinct community structure between tumor suppressive and invasive cancer cell sub-populations. We then showed how ACC may help elucidate ambiguous sub-population annotations in a publicly available human white adipose tissue data set. Finally, we implement a simulation study to validate BANYAN’s ability to recover true community connectivity structure in HST data.

## 1 Introduction

The advent of spatial transcriptomics has allowed for the unprecedented characterization of tissue architecture in terms of spatially resolved transcript abundance [7]. In particular, *high throughput spatial transcriptomics* (HST) technologies such as the 10X Visium platform have become popular due to their deeper transcriptome-wide sequencing depth. The proliferation of HST data has lead to the development of several computational tools for discerning cell sub-populations in HST data, while considering both gene expression and spatial information. The existing tools span a range of methodological categories, including neural networks [12, 22, 11], graph clustering algorithms [13, 19, 31], and Bayesian statistical models [41, 5].

These methods are fundamentally limited in that they do not explicitly model the interactive nature of cell sub-populations in a tissue sample [9]. In other words, the sub-populations derived from existing methods are considered static, and no information is provided on how they relate to one another. Meanwhile, it is known that communication within and between groups of cells is a fundamental driver of healthy and diseased processes in a complex tissue [6]. Moreover, [11] report substantial heterogeneity within traditional mouse olfactory bulb layer annotations, driven in part by spatial variation in intercellular communication patterns. However, detecting higher resolution cell sup-populations with existing tools is challenging as there is no principled methodology for determining which cell sup-populations may be members of a common broader phenotype (e.g., immune or cancer cell sub-types) based on similar yet distinct gene expression or spatial location patterns. As a consequence, current tools cannot be used to study the *community connectivity structure* of cell sub-populations, i.e., the relative similarity among cells within and between subpopulations.

By studying community connectivity structure in HST data, we may obtain valuable insights into the interactive dynamics of cell sup-populations in challenging settings such as the tumor microenvironment. For example, instead of simply labeling categories of immune cells and cancer cells in a tumor, we can describe how these important cell sub-populations relate to one another, and how tertiary intermediate sub-populations may be mediating important dynamics within the tumor microenvironment. Furthermore, characterizing community connectivity structure may help inform more biologically informative annotations of ambiguous sub-populations by relating them to more clearly defined sub-populations. Doing so may allow for a more holistic interpretation of all HST cell clusters in the common case when only a few cell clusters correspond clearly to a known cell type.

To address these gaps, we propose BANYAN (**B**ayesian **AN**alysis of communit**Y** connectivity in sp**A**tial single-cell **N**etworks): a statistical network model capable of discerning community connectivity structure in HST data. BANYAN draws inspiration from the vast field of biological network analysis [17], and is built on the supposition that HST data is most accurately represented as similarity networks that reflect similarity between cell spots in terms of spatial location and transcriptional profiles. To this end, BANYAN introduces a Bayesian multi-layer stochastic block model [28, 36] that infers sub-populations based jointly on transcriptional and spatial similarity between cell spots. This community connectivity structure may then be used to characterize the relationships between cell spots both within and between sub-populations. We offer convenient implementation and interactive visualization functionality via the `R` package `banyan`.

## 2 Results

### 2.1 BANYAN allows for the analysis of community connectivity in HST data

BANYAN is the first HST computational tool to allow for *analysis of community connectivity* (ACC), i.e., the process of inferring the similarity of cell spots within and between sub-populations. A graphical representation is given in Figure 1, and the workflow to achieve ACC can be summarized as follows. First, given cell spot-level gene expression features and spatial coordinate data from HST platforms, we construct two spot-spot nearest neighbors networks. These networks are then integrated into a multi-layer graph data structure. Then, we fit a Bayesian multi-layer stochastic block model (MLSBM), which assumes that spatial location and gene expression patterns of cell spots arise from a common community structure. The estimated parameters from this model allow us to (i) characterize community structure using cell spot sub-population labels, (ii) quantify uncertainty in predicted sub-population labels to identify ambiguous community structure regions, and (iii) infer the community structure of the tissue sample by quantifying the relative similarity between cell spots within and between sub-populations.

**Figure 1:**
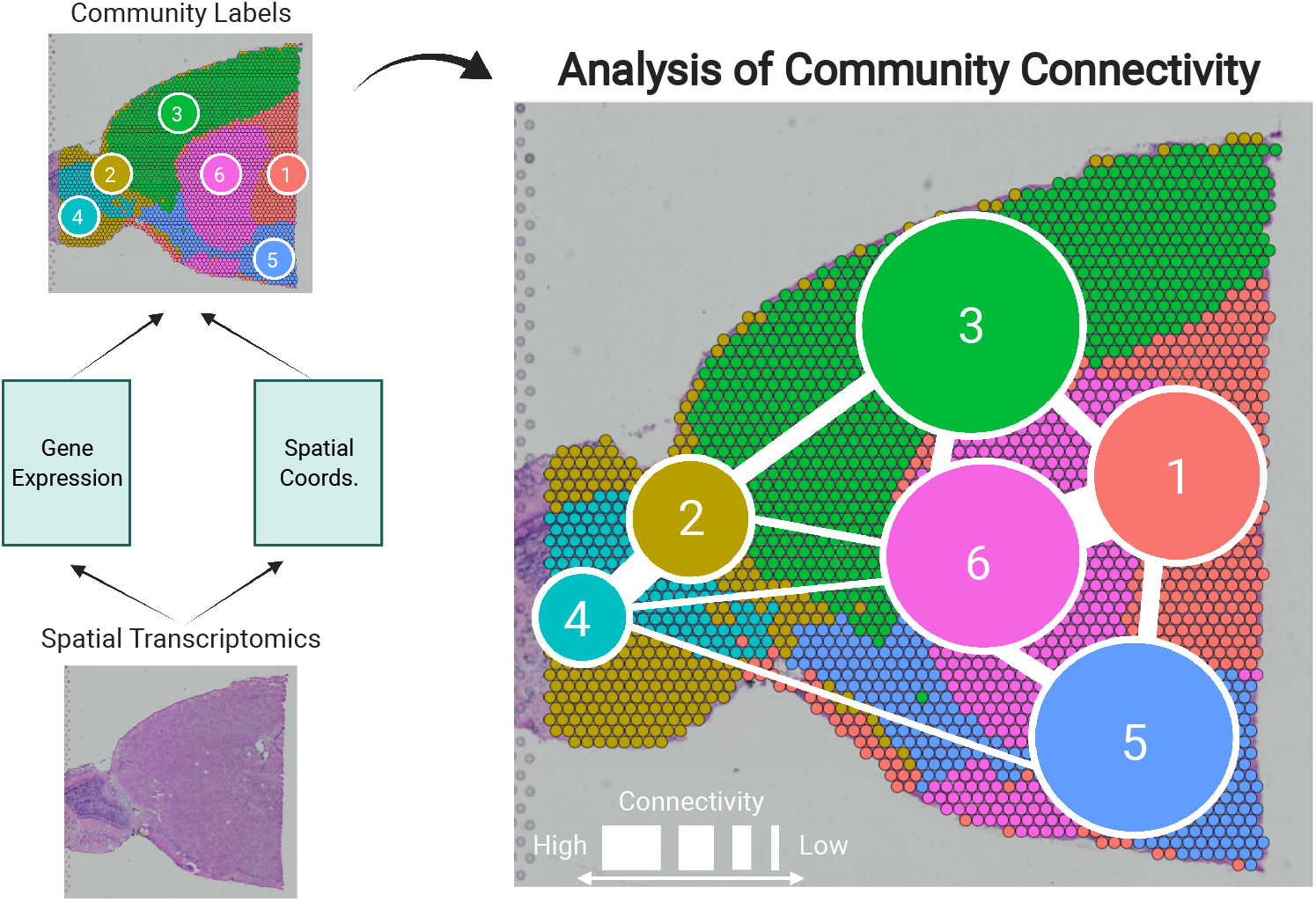
Analysis of community connectivity. Spatial transcriptomics platforms yield gene expression and spatial coordinate matrices, which may be used to derive community labels. Analysis of community connectivity is achieved using BANYAN, which integrates gene expression profiles with spatial locations to infer the connectivity within and between communities.

We provide the `R` package `banyan` for convenient implementation of the proposed workflow. The `banyan` package efficiently implements Bayesian estimation using custom Gibbs sampling algorithms implemented in `C++` using `Rcpp`. The core model fitting functions integrate seamlessly with standard Seurat [35] data structures, allowing users to easily incorporate ACC into existing HST analysis workflows. We have also developed interactive and static visualization functions for interrogation of BANYAN sub-population labels, uncertainty measures, and community connectivity structure. The `banyan` package interfaces seamlessly with standard `Seurat` workflows, and is freely available at https://github.com/carter-allen/banyan.

### 2.2 Discovering community structure in invasive ductal carcinoma

Accounting for roughly 25% of all non-dermal cancers in women, breast cancer ranks as the most common non-dermal female-specific cancer type, and narrowly the most common cancer type across both sexes [39]. Of all sub-types, invasive ductal carcinoma (IDC) is the most common and most severe, accounting for roughly 80% of all breast cancers in women [20]. While previous authors have used spatial trancriptomics to study IDC samples relative to ductal carcinoma in situ (DCIS) samples [40], IDC has yet to be studied through the lens of community structure due to the lack of computational tools available for performing ACC with HST data.

To illustrate ACC in the tumor microenvironment, we applied BANYAN to a publicly available IDC sample sequenced with the 10X Visium platform [4]. We identified five spatially distinct cell spot sub-populations (Figure 2A), with associated uncertainty measures (Figure 2B). We then identified community structure by computing posterior estimates of within and between-community connectivity parameters, displayed in Figures 2C and 2D, respectively. Finally, to interpret each sub-population in terms of IDC biology, we computed the most differentially expressed genes between each sub-population and all others using the Wilcoxon rank-sum test (Figure 2E).

**Figure 2:**
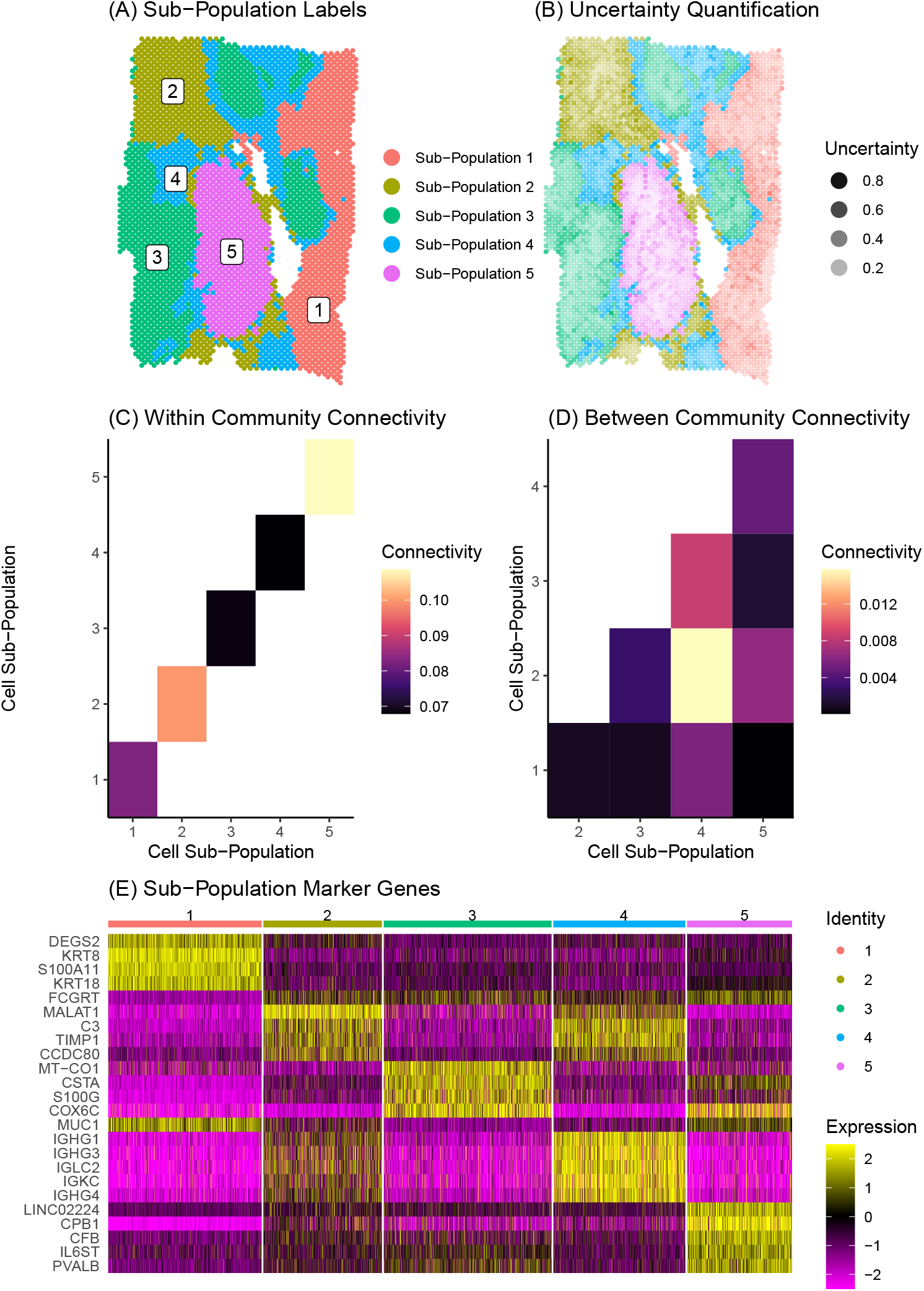
Community structure in invasive ductal carcinoma. (A) Inferred cell spot subpopulation labels from BANYAN. (B) Relative uncertainty measures distinguish uncertain (dark) from certain (light) labels. (C) Within-community connectivity parameters reflect the homogeneity of sub-populations. Higher connectivity values reflect higher homogeneity within sub-populations. (D) Between-community connectivity parameters reflect the relative similarity of cell spots between sub-populations. Higher connectivity values reflect more similarity between sub-populations. (E) Normalized expression of differentially expressed sub-population markers genes.

Figure 2E displays a clear block structure in the expression of sub-population marker genes, indicative of strong community structure signal in the data. These marker genes can be used to obtain a number of interesting biological insights regarding the community structure of the IDC sample. For instance, the *S100A11* gene, a marker for sub-population 1, has been shown to be a diagnostic marker in breast cancers [26] and has been implicated in aggressive tumor progression [27]. Further, *KRT8* is used to differentiate aggressive grades of IDCs [38]. While outside of the context of IDCs, *DEGS2* has been shown to play a role in the invasion and metastasis of colorectal cancer [16]. Taken together, these marker genes suggest sub-population 1 contains a relatively high abundance of aggressive and invasive cancer cell types. On the other hand, sub-population 2 featured marker genes such as *MALAT1* that are associated with tumor suppressive behaviors in IDCs [24]. Another marker gene for sub-population 2, *CCDC80*, has been linked with tumor suppressive functions, albeit not in the context of IDCs [14].

Given these brief characterizations of sub-populations 1 and 2 available from the existing literature, we may hypothesize that these groups of cell spots are in some sense opposed in terms of their role within the tumor based on their transcriptional profiles. Indeed, these sub-populations also reside spatially at opposite ends of the tumor slice. We may investigate the similarity or dissimilarity of these sub-populations 1 and 2 using the between-community connectivity parameters presented in Figure 2D. We find that the estimate of this parameter is near zero (as evidenced by the black coloring of the entry (1,2) in Figure 2D), supporting our hypothesized dissimilarity between sub-populations 1 and 2.

In fact, sub-population 1 featured very low between-community connectivity with all other sub-populations besides sub-population 4, which occupies a heterogeneous “background” position in the spatial landscape of the tissue sample (Figure 2A) and therefore featured relatively high connectivity with all other communities. This spatial heterogeneity is accompanied by relatively low within-community connectivity (Figure 2C), which indicates that spot-spot similarities are less common between cell spots in sub-population 4 than in other sub-populations. In Figure 2E, it can be seen that many of the marker genes for sub-population 2 are shared by sub-population 4, including *MALAT1,* suggesting a similarity between these two sub-populations in terms of transcriptional profiles. In addition to the marker genes shared with sub-population 2, sub-population 4 features several of its own distinct marker genes, namely the immunoglobulin heavy chain-encoding RNAs *IGHG1* and *IGHG3.* These genes have themselves been shown to feature tumor suppressive tendencies via promotion of B cell specific immunoglobulin [21], and have been associated with increased patient survival [25]. This observation of functional similarity between sub-populations 2 and 4 is validated by Figure 2D, which clearly shows the highest estimated between-community connectivity in the data occurring between sub-populations 2 and 4. Taken together, these observations may lead us to reason that the sub-population 1 vs. 2 dynamic described previously is linked via the more heterogeneous yet still tumor suppressive-like sub-population 4. While these observations would require further experimental validation to confirm, they showcase the unique ability of BANYAN to describe community structure in the data.

### 2.3 Characterizing ambiguous annotations in human white adipose tissue

Next, to demonstrate the application of ACC to inform ambiguous sub-populations, we applied BANYAN to the analysis of a human white adipose tissue (WAT) sample sequenced with the 10X Visium platform [8]. In contrast to the IDC data set considered in Section 2.2, WAT samples are characterized by weak histological organization, thus challenging the manual annotation of cell spots. In Figure 3A, we display the manual annotations from [8] for an individual WAT sample (ID: ADI24). Of the 2,747 cell spots in the original sample, 6 were annotated “unknown” and 1,520 were labeled “unspecific.” Hence, over 50% of cell spots were unable to be clearly annotated due to ambiguous expression profiles or heterogeneous cell type mixtures within cell spots as a result of the resolution of the 10X Visium platform, a matter further complicated by the weak histological organization of WAT samples. Since existing approaches to labeling sub-populations in HST data fail to account for the community structure of tissue samples, disambiguating unknown cell spots *post hoc* remains challenging with existing tools.

**Figure 3:**
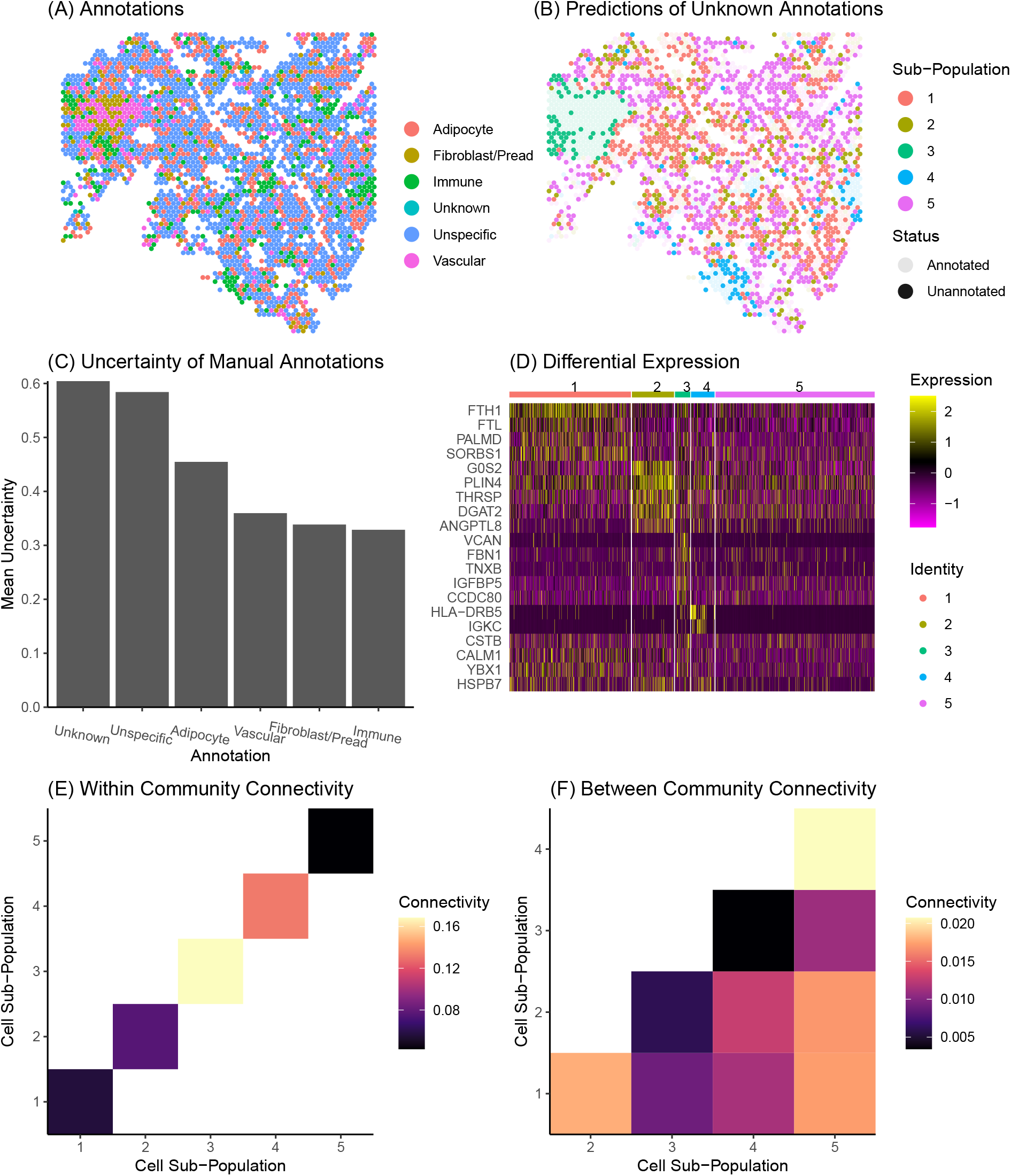
Characterization of ambiguous cell spot annotations. (A) Manual annotations. (B) BANYAN labels of unknown or unspecific (i.e., unannotated) cell spots from (A). (C) Mean BANYAN uncertainty scores by each annotation group. (D) Differential expression of BANYAN sub-populations using the unannotated subset. (E) Within-community connectivity between cell spots belonging to each sub-population. (F) Between-community connectivity between cell subpopulations.

In Figure 3B, we show the sub-population labels for the entire WAT sample ADI24 derived from BANYAN, where the unannotated cell spots (i.e., those classified as either “unknown” or “unspecific” by [8]) are highlighted in bold. We find that BANYAN identified residual heterogeneity within the 1,526 unannotated cell spots, with 513 unannotated cell spots labeled as sub-population 1, 178 unannotated cell spots labeled as sub-population 2, 64 unannotated cell spots labeled as sub-population 3, 99 unannotated cell spots labeled as sub-population 4, and 672 unannotated cell spots labeled as sub-population 5. We quantified the average uncertainty measures derived from BANYAN across the original annotation groups, and found the highest uncertainty occurring in the “unknown” or “unspecific” categories (Figure 3C), further validating the low signal contained in this subset of cell spots.

In Figure 3D, we present a heatmap depicting differentially expressed marker genes for each BANYAN sub-population using only the unannotated cell spots. The results from this analysis suggest that residual heterogeneity exists within the unannotated cell spot subset, with distinct marker genes present for sub-populations 1 through 4. Meanwhile, BANYAN sub-population 5 lacked clear markers, suggesting this sub-population could be reflective of a smaller ambiguous subset of cell spots within the original unannotated subset. In particular, as shown in the within-community structure displayed in Figure 3E, we find sub-population 3 to exhibit the highest within-community connectivity. This high within-community connectivity is supported by the contiguous spatial organization of sub-population 3 (Figure 3B) as well as high expression of distinct marker genes such as *VCAN* (Figure S1), which are suggestive of veriscan producing adipocytes associated with the development obesity-related inflammation of adipose tissue [18].

When assessing the between-community structure of the WAT sample inferred by BANYAN as shown in Figure 3F, we find additional evidence for sub-population 5 being a heterogeneous unspecific sub-population (e.g., low within-community connectivity in Figure 3E and high between-community connectivity in Figure 3F). However, sub-population 5 did feature relatively high connectivity with sub-population 4, as evidenced by the bright coloring of entry (4,5) of Figure 3F. Sub-population 4 was marked by significant differential expression of adipose-resident immune cell related genes (Figure S1) such as *IGKC* [37]. This suggests sub-population 5 may play an important role in mediating the function of immune cells within body fat compartments [37]. While additional studies correlating these sub-populations with true single-cell data such as single cell RNA-seq would aid in further elucidation of these ambiguous sub-populations, leveraging community structure allows for more informative interpretation even in low-resolution HST data.

### 2.4 Simulation studies show BANYAN identifies community structure in a broad range of signal-to-noise ratio settings

Finally, we designed a simulation study to validate the performance of the MLSBM employed by BANYAN, in the sense of identifying cell sub-populations and recovering community connectivity structure. We adopted a publicly available sagittal mouse brain data set [3] sequenced with the 10X Visium platform. We manually allocated the *N* = 2696 total cell spots in the original sagital mouse brain data set into one of *K* = 4 simulated ground truth tissue segments, resulting in 4 spatially contiguous mouse brain layers (Figure 4A). The result of this was a ground-truth community structure that is reflective of sub-populations found in real HST data. We then simulated the spot-spot gene expression similarity network from a stochastic block model with community structure given by

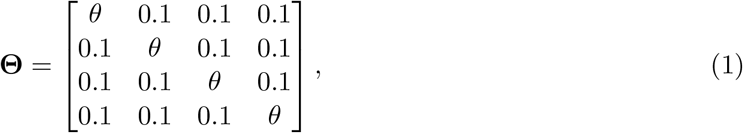

where the *signal to noise ratio* (SNR) of the simulated gene expression network is given by SNR = *θ*/0.1. SNR values much greater than 1 give rise to a strong community structure in the simulated data, while SNR values close to 1 result in a weaker community structure. We do not consider values of SNR below 1, as the resultant dissortative community structure is not reflective of cell type structure in HST data. We simulated gene expression networks for a range of SNR settings using *θ* = (0.105, 0.11, 0.13, 0.15, 0.17, 0.20, 0.22, 0.25) and fit two model variants: (i) a single-layer model considering gene expression information only, and (ii) the full MLSBM using both gene expression and spatial networks.

**Figure 4:**
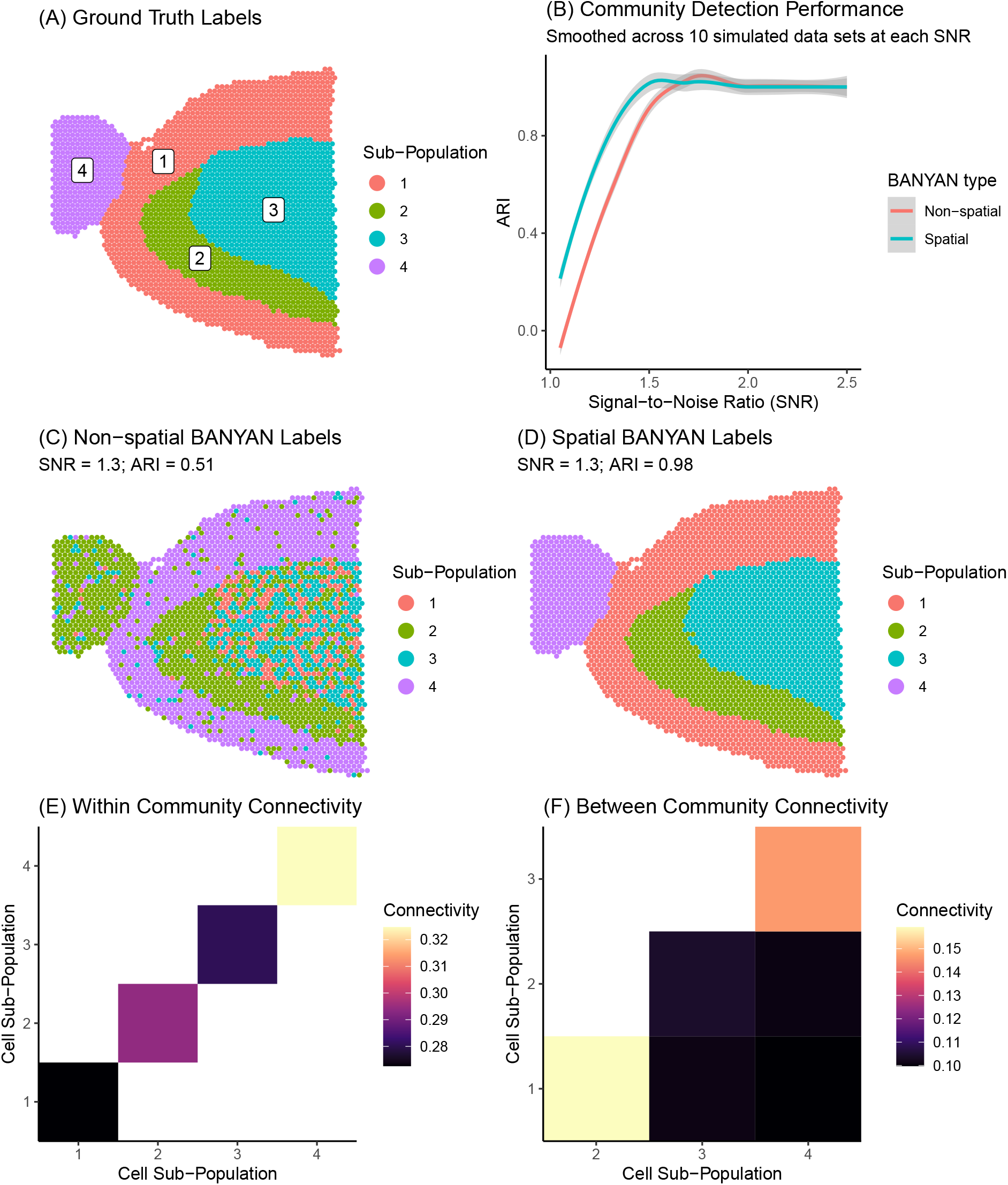
Segmenting sagittal mouse brain tissue sample to four different clusters. (A) Ground truth labels. (B) Average adjusted rand index (ARI) vs. signal-to-noise ratio (SNR). (C) Estimated BANYAN labels from the simplified single-layer non-spatial model. (D) Estimated BANYAN labels from multi-layer spatial model. (E) Within-community connectivity estimates. (F) Between-community connectivity estimates.

Figure 4B displays the average adjusted rand index (ARI) – a measure of accuracy in cell spot labels relative to the ground truth labels in Figure 4A, for the single-layer non-spatial approach and the multi-layer spatial approach. We find that at low SNR settings (e.g., below 1.5) the spatial model outperforms the non-spatial model in recovering ground truth cell spot labels. This is indicative of BANYAN’s ability to leverage spatial information to detect community structure in low-signal data. At higher SNR settings, the strong community structure signal contained in the simulated gene expression layer is sufficient for accurate community structure recovery, and the spatial information does not provide any further information.

We showcase the community structure results for one particular simulated data set at a setting of SNR = 1.3, reflective of a relatively low signal setting. Figure 4C displays the inferred community structure from the single-layer non-spatial model, while Figure 4D shows the same for the multilayer spatial model. We find that at this moderately low signal setting, the non-spatial model is unable to accurately recover true cell spot labels, while the spatial model predicts cell spot labels almost perfectly. These results are indicative of the ability for spatial information to aid in disambiguating community structure in low-signal data settings.

While the simulated HST data generated from the community structure encoded in Equation (1) features a homogeneous community structure (i.e., uniform within and between-community connectivity parameters), BANYAN is capable of detecting more heterogeneous community structures.

To illustrate this, we generated a simulated mouse brain HST data set from

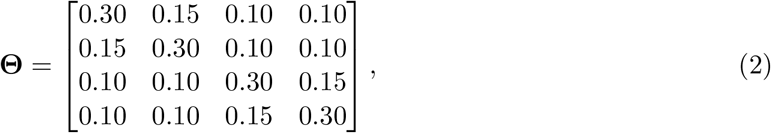

which features 50% stronger connectivity between sub-population pairs (1,2) and (3,4) than other between-community pairs. This setting reflects the common real data scenario wherein cliques of sub-populations form. In Figures 4E and 4F, we display the estimated within and between-community connectivity parameters, respectively. We find that the BANYAN model correctly identified the sub-population cliques (1,2) and (3,4) as sharing higher between-community connectivity than the remaining sub-population pairs, showcasing the model’s ability to identify heterogeneous community structures. In Figure S2 of the Supplementary Materials, we demonstrate how the Bayesian information criterion (BIC) identifies the true number of communities in the simulated data, validating the use of statistical model fit criteria for choosing the number of sub-populations in the absence of prior knowledge.

## 3 Discussion

We have proposed BANYAN: a network-based statistical framework for analysis of community connectivity in HST data. We applied BANYAN to human breast cancer and white adipose tissue to illustrate its utility in applied settings. In the breast cancer case study, we found a strong community structure, with sub-populations marked by both invasive cancer and cancer suppressive marker genes. Using community structure parameters, we also identified an intermediate subpopulation between these two. In the white adipose tissue case study, we demonstrated the use of BANYAN to disambiguate unknown or unspecific cell spot labels. In our simulation study, we validate BANYAN’s ability to accurately identify the tissue architecture, especially in the low signal setting, and to recover the community connectivity structure.

There a number of ways our work may be extended. First, often the SBM is refined to accommodate heterogeneous degree distributions among nodes, i.e., *degree correction* [23]. By making this methodological extension to the MLSBM at the core of BANYAN, one could relax our assumption that each cell spot features the same number of neighbors and thereby allow for certain cells spots to feature more connections to the rest of the tissue than other cell spots, such as those on the periphery of the tissue sample. Learning the degree of each cell spot would then inform the detection of highly connected “hub” regions, or weakly connected “satellite” regions of a tissue sample. Second, the inherent complexity of network data structures leads to a heavy computational burden for large HST experiments. Another extension could be to relax the assumption that gene expression and spatial location layers are governed by common community structure parameters, instead allowing for layer-specific interpretations of community structure. While we implement our proposed Markov chain Monte Carlo (MCMC) sampling algorithm using efficient `Rcpp` routines, BANYAN still requires significantly more computational time than non-network statistical methods [5, 41]. Further optimization would help to reduce computational burden of community connectivity analysis. Finally, while BANYAN provides the first statistical framework for quantifying community connectivity structure in HST data, further extensions could be made to link BANYAN with methods for predicting cell-cell interactions using data such as ligand-receptor pair status of cells. By doing so, one could refine the general notion of cell spot connectivity to cell spot interaction, which is of major interest in HST data analysis. In this sense, BANYAN establishes a promising statistical framework that may be extended to a wide range of analyses focused on investigating the interactive nature of HST data.

## 4 Methods

### 4.1 Data pre-processing

To represent the interactive nature of cells and cell types, we adopt two cell-cell similarity networks as our primary data objects: one for gene expression and another for spatial location. To form the cell spot-cell spot gene expression similarity network, we first apply standard pre-processing steps including scaling, removal of technical artifacts, and identification of highly variable genes [19, 2, 1]. We then embed each of the *N* total cell spots in a lower-dimensional space using principal components analysis (PCA) applied to the top 2,000 most variable genes. To form the cell spot-cell spot gene expression similarity matrix, we represent each cell spot as a node and connect each cell spot to its *R* closest neighboring cell spots in the gene expression principal component space using a binary edge. We utilize the same approach to construct the spatial cell spot-cell spot similarity network, where principal components are replaced with 2-dimensional spatial coordinates. The resultant data structure is two networks with *N* nodes, each of degree *R*. By default, we adopt the widely used heuristic of choosing *R* as the closet odd integer to 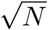 [34], which allows the number of neighboring spots to increase as the size of the tissue sample increases. With the typical HST experiment yielding a total number of cell spots between 2,000 and 3,000, this heuristic leads to consideration of between third and fourth order neighborhood structures (Fig 5). Overall, we view *R* as a tuning parameter that may be adjusted depending on the amount of information sharing desired across a tissue sample.

**Figure 5:**
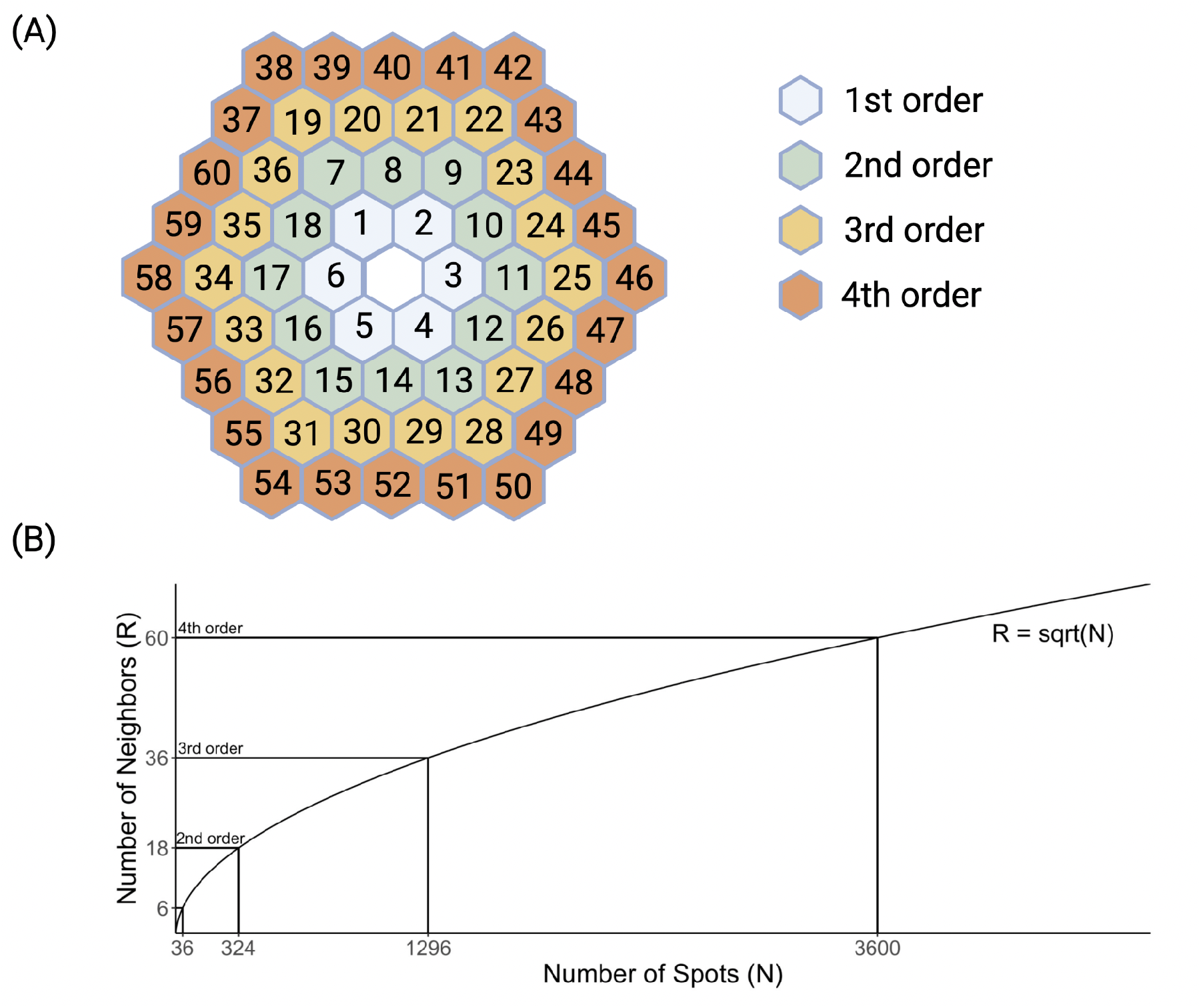
Graphical depiction of relationship between number of neighbors and neighbor order. (A) Hexagonal neighborhood structure for an interior cell spot shown with 1st through 4th order neighbors. (B) Suggested relationship between the number of cell spots (N) and the number of nearest neighbors (R).

### 4.2 Model

We develop the core statistical model within BANYAN as an extension of the widely used stochastic block model (SBM) [33], a flexible generative model for network data that allows for the assessment of community structure based on the frequency of binary edges among and between subsets of nodes. We define **A**^1^ as the *N* × *N* binary adjacency matrix encoding the gene expression similarity network, and **A**^2^ as the binary adjacency matrix encoding the spatial similarity network. The matrix elements 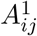 and 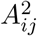 indicate the presence or absence of a binary un-directed edge between nodes *i* and *j* for gene expression and spatial information, respectively. We define 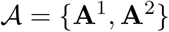 as the multi-layer graph that encodes similarity between cell spots in terms of both gene expression and spatial information. While we focus on integration of spatial and gene expression information, our proposed framework may be extended to *L* layers to incorporate other sources of information from multiplexed experimental assays.

Given the multi-layer graph data 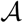, we assume that the absence or presence of edges in each layer between each pair of nodes *i* and *j* follows a Bernoulli distribution with probability of an edge *θ_zi,zj_*, where *z_i_* ∈ {1,…,*K*} denotes the latent cell spot sub-population assignment for cell spot *i*. We refer to such a model as a multi-layer stochastic block model (MLSBM). Formally, we assume for *l* = 1, 2,

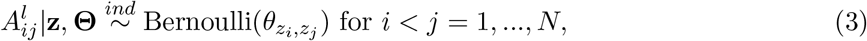

where **z** = (*z*_1_,…,*z_N_*), and **Θ** is a *K* × *K connectivity matrix* with diagonal elements *θ_rs_* for *r* = *s* = 1,…, *K* controlling the probability of an edge occurring between two cell spots in the same sub-population, and off-diagonal elements *θ_rs_* for *r* < *s* = 1, …,*K* controlling the probability of an edge occurring between two nodes in different cell spot sub-populations. Importantly, Model (3) implies that connections among cell spots in the gene expression and spatial layers are governed by a common set of community structure parameters **z** and **Θ**. Given Model (3) and data 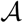, our primary inferential objective is to characterize cell spot sub-populations and the cell-cell interaction both *within* and *between* them by estimating the parameters **z** and **Θ**, which we accomplish using a Bayesian approach as described below.

### 4.3 Bayesian Inference

#### 4.3.1 Priors

To achieve a fully Bayesian parameter estimation scheme, we assign prior distributions to all model parameters. We adopt available conjugate priors to obtain closed-form full conditional distributions of all model parameters, allowing for straightforward Gibbs sampling. For the latent cell sub-population indicators *z*_1_,…,*z_N_*, we assume a conjugate multinomial-Dirichlet prior with 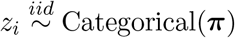 for *i* = 1,…,*N*, and **π** ~ Dirichlet(*α*_1_,…,*α_K_*), where **π** = (*π*_1_,…,*π_κ_*) controls the relative size of each cell sub-population to allow for a heterogeneous distribution of cell type abundances. We adopt a conjugate Beta-Bernoulli prior for **Θ** by assuming 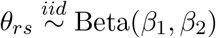 for *r* < *s* = 1,…*K*. As a default, we opt for weakly informative priors by setting *α*_1_ = *α*_2_ = … = *α_K_* = 1 and *β*_1_ = *β*_2_ = 1 [15].

#### 4.3.2 Markov chain Monte Carlo (MCMC) algorithm

The model proposed in Sections 4.2 and 4.3.1 allows for closed-form full conditional distributions of all model parameters. Thus, we adopt the following Gibbs sampling algorithm for parameter estimation. In practice, we recommend initializing the indicators *z_i_*,…,*z_N_* using a heuristic graph clustering method such as the Louvain algorithm [10] applied to **A**^1^ to facilitate timely model convergence.

1. Update **π** from its full conditional (**π**| **A**, **z**, **Θ**) ~ Dirichlet(*a*_1_,…,*a_N_*), where *α_k_* = *α_k_*+ *n_k_*, and *n_k_* is the number of nodes assigned to cell sub-population *k* at the current MCMC iteration, i.e., 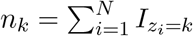.
2. For *r* ≤ *s* = 1,…, *K*, update *θ_rs_* from

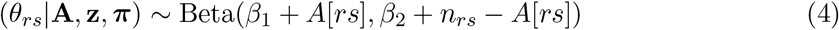

where *A*[*rs*] are the number of observed edges between communities *r* and *s* across both layers, and *n_rs_* = 2(*n_r_n_s_* – *n_r_I*(*r* = *s*)) are the number of possible edges between communities *r* and *s, n_r_* is the number of nodes assigned to cell sup-population *r*, and *I*(*r* = *s*) is the indicator function equal to 1 if *r* = *s* and 0 otherwise.
3. For *i* = 1,…,*N*, update *z_i_* from (*z_i_*|*z_–i_*, **A, π, Θ**) ~ Categorical(***ρ***_*i*_), where *ρ*_*i*1_ = (*ρ*_*i*1_,…,*ρ_iK_*) and

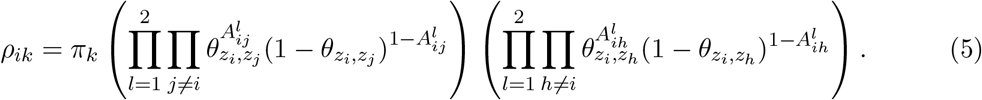

#### 4.3.3 Model selection

The choice of number of cell sub-populations *K* is a critical step in the analysis of HST data. In some cases, *K* may be chosen based on *a priori* biological knowledge of the cell types expected to exist in a tissue sample, or the desire to investigate a known number of sub-populations within a more homogeneous tissue sample. In the absence of such prior information, *K* may be chosen using statistical model fit criteria, such as the Bayesian information criterion (BIC) [32].

#### 4.3.4 Label switching

Label switching is a ubiquitous issue faced by models whose likelihood is invariant to permutations of a latent categorical variable such as **z**. Consequently, stochastically equivalent permutations of **z** may occur over the course of MCMC sampling, causing the estimates of all community-specific parameters to be conflated, thereby jeopardizing the accuracy of model parameter estimates. Previous approaches for addressing label switching rely on re-shuffling posterior samples after completion of the MCMC algorithm [29]. However, such methods rely on prediction and thereby are subject to to prediction error. To protect against label switching within the MCMC sampler, we adopt the canonical projection of **z** proposed by [30], who restrict updates of **z** to the reduced sample space 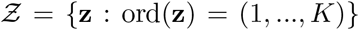, wherein label switching is less likely due to the restricted sample space. In practice, we manually permute **z** at each MCMC iteration such that community 1 appears first in **z**, community 2 appears second in **z**, *et cetera*. Finally, we estimate **z** using the maximum *a posteriori* (MAP) estimate across all post-burn MCMC samples [15].

### 4.4 Analysis of community connectivity

Estimation of the MLSBM model parameters **Θ** and **z** with the corresponding maximum *a posteriori* estimates 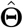 and 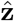 allows for inference of community connectivity structure in HST data. While the estimated community labeling vector 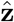 is what we use to define communities, the elements of 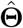 describe how cell spots within and between communities relate to one another, thereby characterizing community connectivity. Specifically, elements 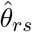 reflect the estimated probability of a randomly chosen cell spot in community *r* sharing a nearest neighbors edge in 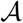 with a cell spot in community *s*. When *r* = *s*, 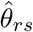 reflects the average connectivity within a community, which may be used to assess the relative homogeneity of a community. Heterogeneous communities tend to have lower average within-community connectivity, while more homogeneous communities tend to have higher within-community connectivity. Likewise, when *r* = *s*, 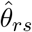 represents the probability of connection between cell spots in two distinct communities. This between-community connectivity measurement allows us to discern closely related communities that may contain similar cell types from more distinct communities. Taken together, these between and within-community connectivity parameters capacitate analysis of community connectivity.

### 4.5 Uncertainty quantification

Discrete clustering approaches for community structure identification inherently fail to account for heterogeneity within each spot cluster by assuming all cell spots within the same cluster are stochastically equivalent. To address these shortcomings of existing approaches, we utilize the inferential benefits of Bayesian modeling to derive two biologically relevant measures: (i) continuous phenotypes and (ii) uncertainty scores. With continuous phenotype measures, we may assess the propensity of a given cell spot for a cell type other than its most likely cell type, thus allowing for identification of possible intermediate cell states. Relatedly, uncertainty measures allow us to distinguish high confidence from low confidence cell type assignments.

Conceptually, we choose *c_ik_*, the continuous phenotype for cell spot *i* towards cell sub-population *k*, to be proportional to the posterior probability *P*(*z_i_* = *k*|*z*_–*i*_, **A, π, Θ**). Considering terms only related to *z_i_*, we define *c_ik_* as

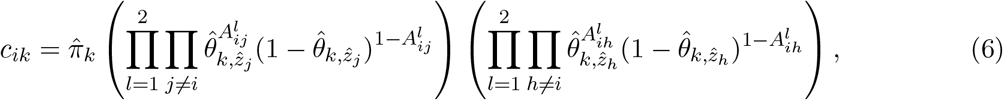

for *i* = 1,…,*N* and *k* = 1,…,*K*, where 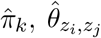, and 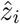 are the posterior estimates of *π_k_, θ_z_i_,z_j__*, and *z_i_*, respectively. We define the uncertainty measure for cell spot *i* as 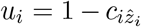, i.e., the sum of cell spot *i* continuous phenotypes for all cell types besides 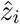.

## Supplementary Information

**Figure S1: Differentially expressed markers for BANYAN sub-populations across all cell spots in white adipose tissue (WAT) sample.** Differential expression p-values were computed using the Wilcoxon rank-sum test. Sub-populations 1 through 4 display clear marker genes while sub-population 5 remained more ambiguous.

**Figure S2: Identification of true number of communities in simulated data using the Bayesian information criterion (BIC).** Higher values indicate better model fit.

## Acknowledgements

This work has been supported through grant support from the National Human Genome Research Institute (R21 HG012482), National Institute of General Medical Sciences (R01 GM122078 and R01 GM131399), National Institute on Drug Abuse (U01 DA045300), National Institute on Aging (U54 AG075931), the National Science Foundation (NSF1945971), and the Pelotonia Institute of Immuno-Oncology (PIIO). The content is solely the responsibility of the authors and does not necessarily represent the official views of the funders.

